# Plate-Q: A Frugal Microplate-Reader for Bacterial Signal Quantification

**DOI:** 10.1101/2022.06.11.495768

**Authors:** Varun Sendilraj, Madhav Gulati, Janet Standeven, Saad Bhamla

**Affiliations:** Bhamla Lab, Georgia Institute of Technology, Atlanta GA

**Keywords:** Microplate Reader, Optical Device, Frugal Science, Open-Source, Computer Vision

## Abstract

Microplate readers are laboratory instruments used to measure biological properties and reactions within a microplate. A microplate consists of small wells in which separated reactions take place. These reactions convert the progression of biochemical processes into optical signals. A microplate reader detects these signals and quantifies a parameter of interest. Most lab-grade microplate readers can cost up to $20,000, making them unaffordable for underfunded labs and developing countries. In light of this, we developed Plate-Q, a $150 frugal microplate reader capable of quantifying green fluorescence protein and optical density of bacterial samples from 96-hole microplates. Rather than using optical sensors found in laboratory microplate readers, Plate-Q takes advantage of a Raspberry-Pi camera to capture images of a microplate and manually extract brightness values using computer vision algorithms. Plate-Q utilizes 440nm excitation lights with a 510nm emission filter to measure GFP expression and 600nm lights for optical density readings. However, camera-based technologies have three main problems: lens distortion, signal interference, and brightness calculations. Plate-Q addresses these problems through camera movement over the well-plate, image optimization, 3D printed well-plate covers and mathematical regression algorithms. In testing with fusarium dual-plasmid biosensors, Plate-Q showed comparable accuracy to commercial microplate readers with an average error of .034 and a root mean squared error of .012 for fluorescence and an average error of .045 and root mean squared error of .023 error for optical density. Users can scan for other fluorescent proteins by changing the light source and filter for different wavelengths. With all aspects of Plate-Q being open source, labs worldwide can quantify samples in an affordable and time-efficient manner.

## Introduction

### Laboratory Grade Microplate Readers

Microplate readers, also known as plate readers or microplate photometers, are instruments used to detect biological, chemical, or physical signals of samples in microplates. This device is widely used in biotechnology research, drug discovery, bioassay validation, quality control, and in manufacturing processes within the pharmaceutical and biotechnological industries. The most common microplate format used in academic research laboratories or clinical diagnostic laboratories is 96-well (8 by 12 matrix) with a reaction volume usually between 100 and 200 μL per well [8]. Detection modes for microplate readers include absorbance, fluorescence intensity, luminescence, time-resolved fluorescence, and fluorescence polarization with the most common being the first two, respectively. A microplate reader is made up of a supercontinuum laser, excitation filter, dichroic mirror, emission filter, and high sensitivity photomultiplier tubes which are all controlled through motorized movement over the microplate. The supercontinuum laser acts as the light source for the samples, which is adjusted to the right excitation wavelength through the excitation filter. After exciting the samples, the dichroic mirror guides the bacterial signals to the emission filter and the high sensitivity photomultiplier tube for quantification [9]. The many parts of laboratory-grade microplate readers constitute a high price of around $15,000 to $20,000, making them unaffordable to underfunded labs and developing countries worldwide.

### Our Solution

We present Plate-Q, a frugal microplate reader for $150 that can quantify the fluorescence intensity of green fluorescent proteins (GFP) and absorbance (optical density) of bacterial samples in a 96-well microplate. In developing Plate-Q, we isolated the most expensive components in a laboratory-grade microplate reader to develop a more low-cost, reproducible solution. The first change is the mechanism of the excitation source. The supercontinuum laser and excitation filter were swapped for an evenly distributed LED light source falling within GFP’s excitation wavelength range of 440 nanometers and 600 nanometers for absorbance. Additionally, we replaced the sensing of the high sensitivity photomultiplier tube with a high sensitivity CMOS camera. The camera utilizes computer vision and mathematical regression algorithms to isolate and quantify the brightness of the samples. With the use of the evenly distributed excitation light over the microplate along with the camera-based quantification, we eliminate the need for motorized movements. Plate-Q is completely open-source and users can customize the design to scan for other fluorescent proteins by replacing the light source and filter for different wavelengths.

## Materials And Design

### Parts List

- Frosted Acrylic Plate
- Raspberry Pi
- Raspberry Pi HQ Camera
- 16mm Focal Length Lens
- 440 nm LED Strip Lights
- 600 nm LED Strip Lights
- 510 nm Glass Emission Filter

### Past Design

The design of Plate-Q has been revised many times. The initial design of Plate-Q contained a single focal light source that would not evenly distribute light across the well plate and used angle and distance characterization to account for differences in light intensity caused by the uneven light distribution (Figures. 1 & 2). After thorough testing, we quickly learned that characterizing for angle and distance is impractical and yields inaccurate results.

**Figure 1.**
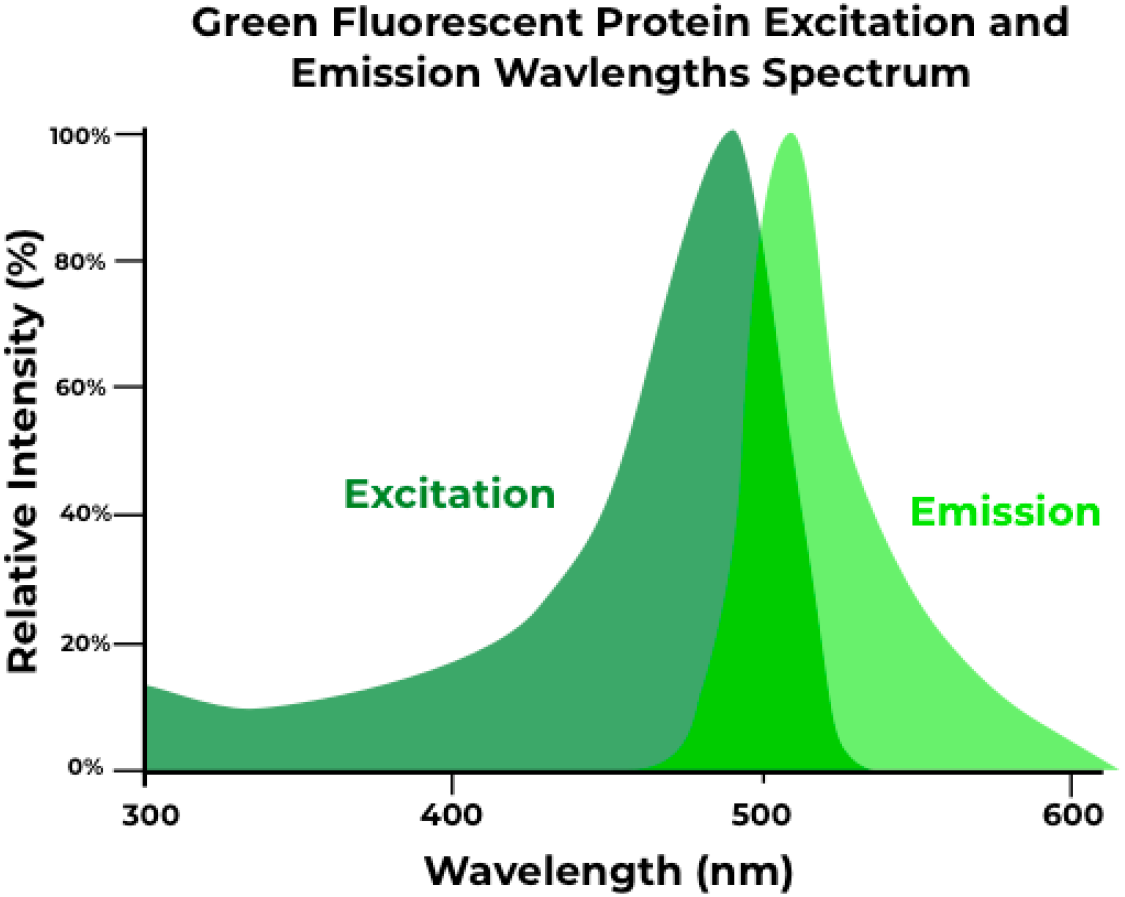
Emission and Excitation Spectrums of Green Fluorescent Protein.

**Figure 2.**
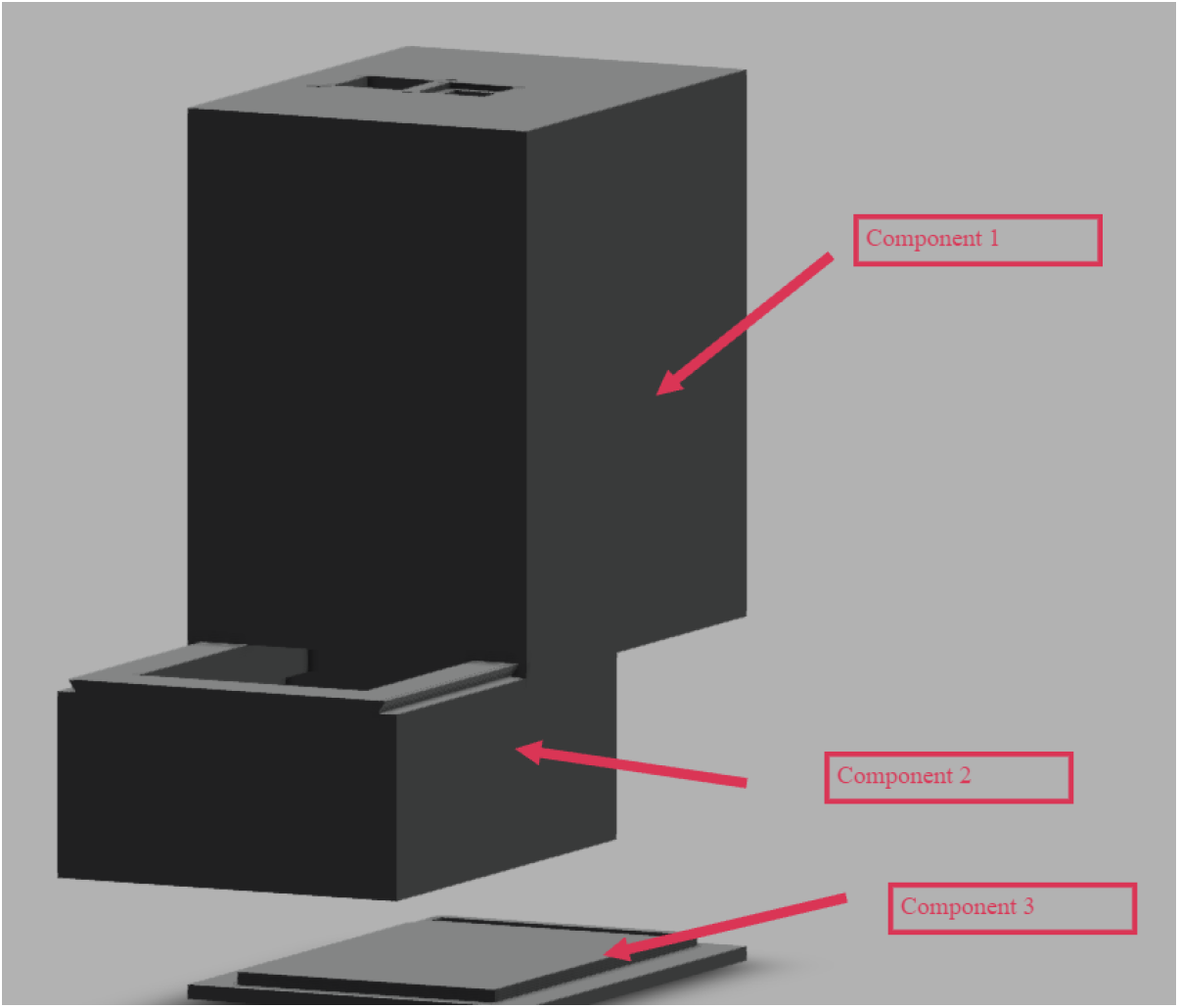
Original Plate Reader Design with Light Sources housed on top without even light distribution on the sample.

**Figure 3.**
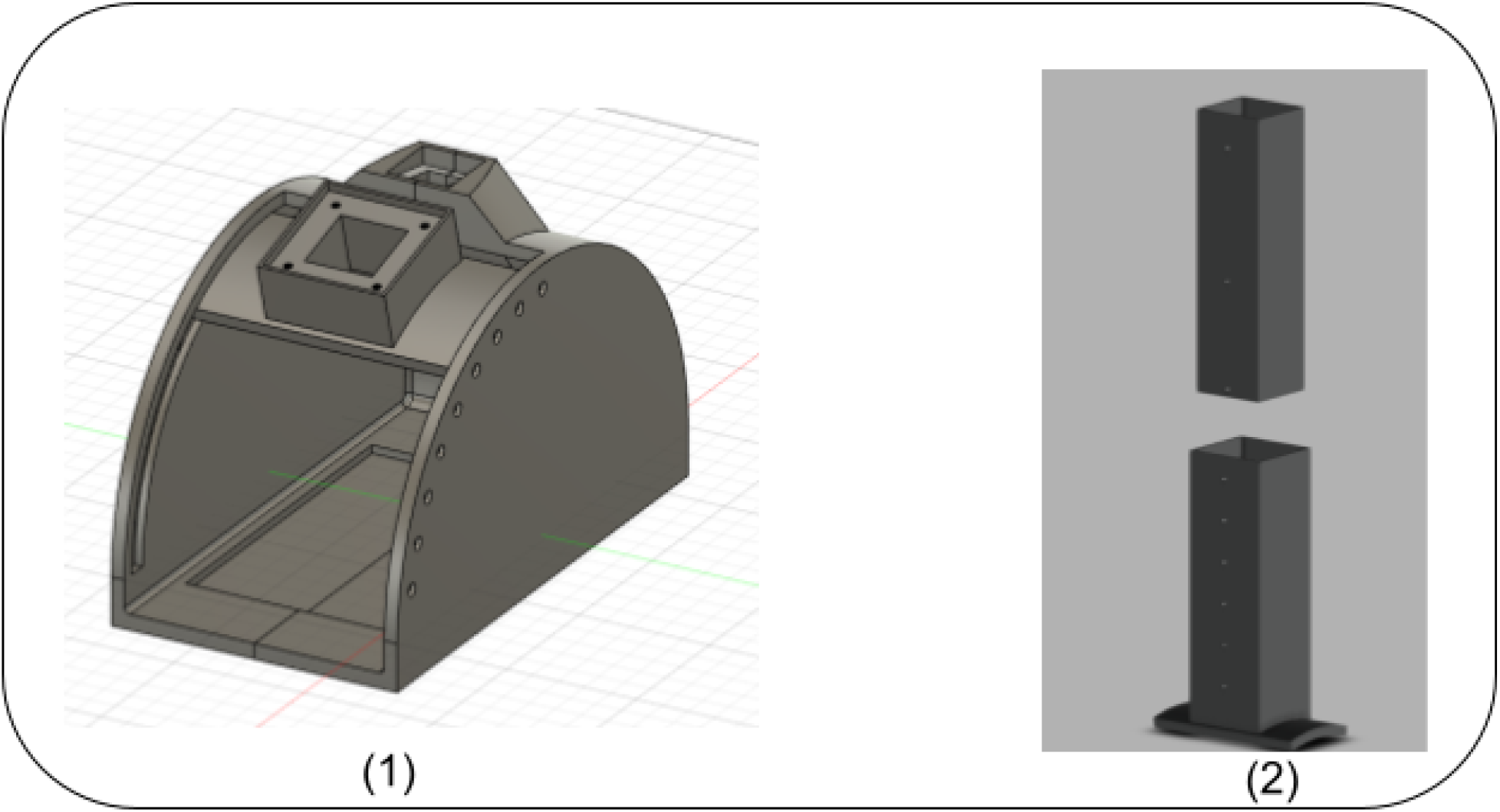
3-1. Testing jig of Plate-Q that revolved its light source around a center of rotation to simulate multiple angles of excitation. 3-2. A telescopic arm that attaches to the light source to simulate different distances.

### Current Design

Plate-Q is broken down into 3 detachable parts: Top cover (Figures. 4-1), a Light-diffusing chamber (Figures. 4-2), and a bottom cap (Figures. 4-3). The Raspberry Pi camera is housed on the top cover and has 4 positions of movement. The cover is printed with a matte black filament which helps prevent the reflection of light into the camera. This part then slides onto the light-diffusing chamber. The 96-well plate is placed into the slot on top of the light-diffusing chamber (Figures. 5). We use a frosted acrylic diffuser that is able to diffuse the light from the LED strips evenly across the well plate(Figures. 6). The bottom cap, which houses the led light strips, will be under the diffuser in order to provide the GFP and OD600 wavelengths needed for quantification.

**Figure 4.**
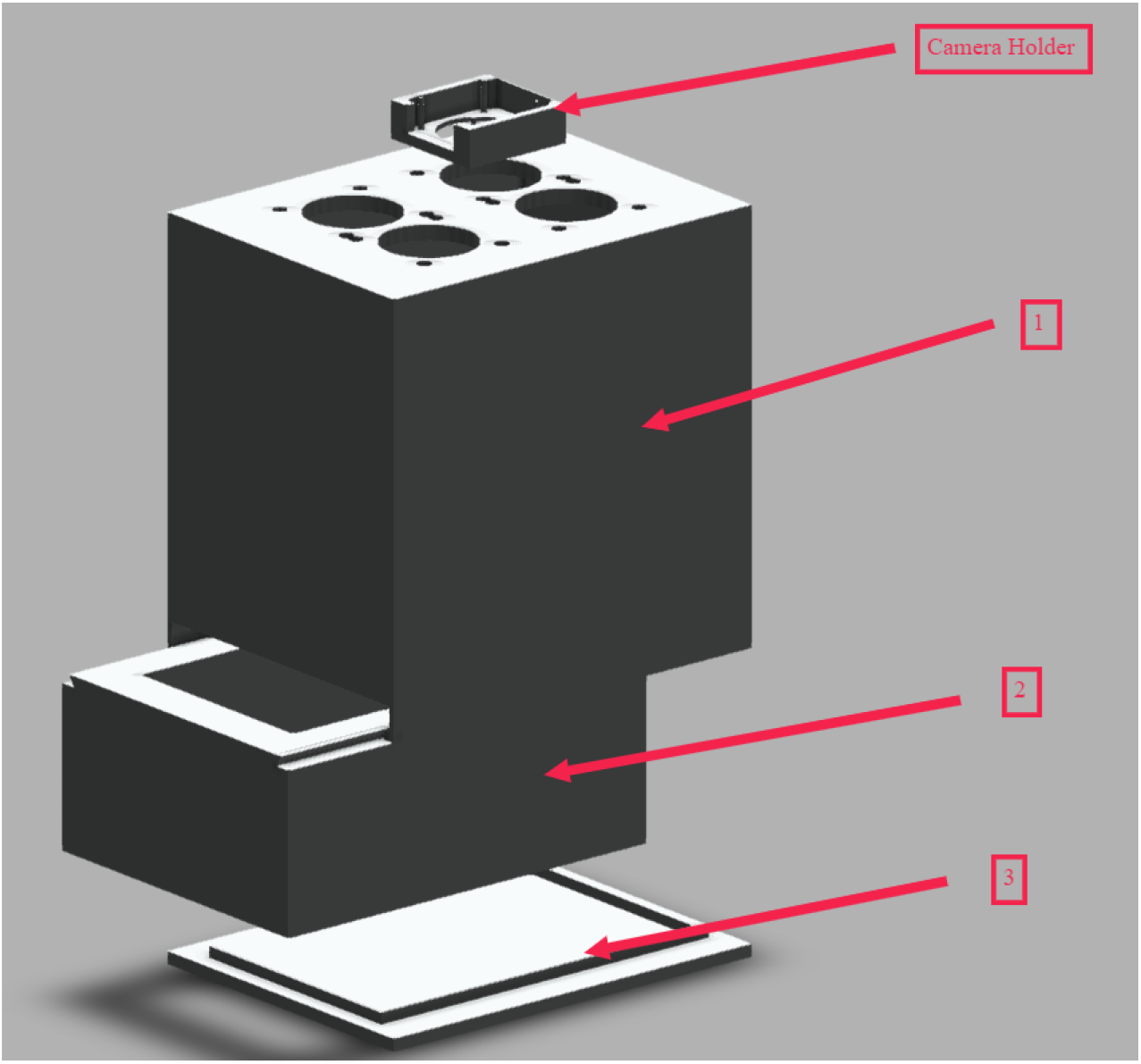
Current design of Plate-Q with even light distribution and camera movement over the microplate

**Figure 5.**
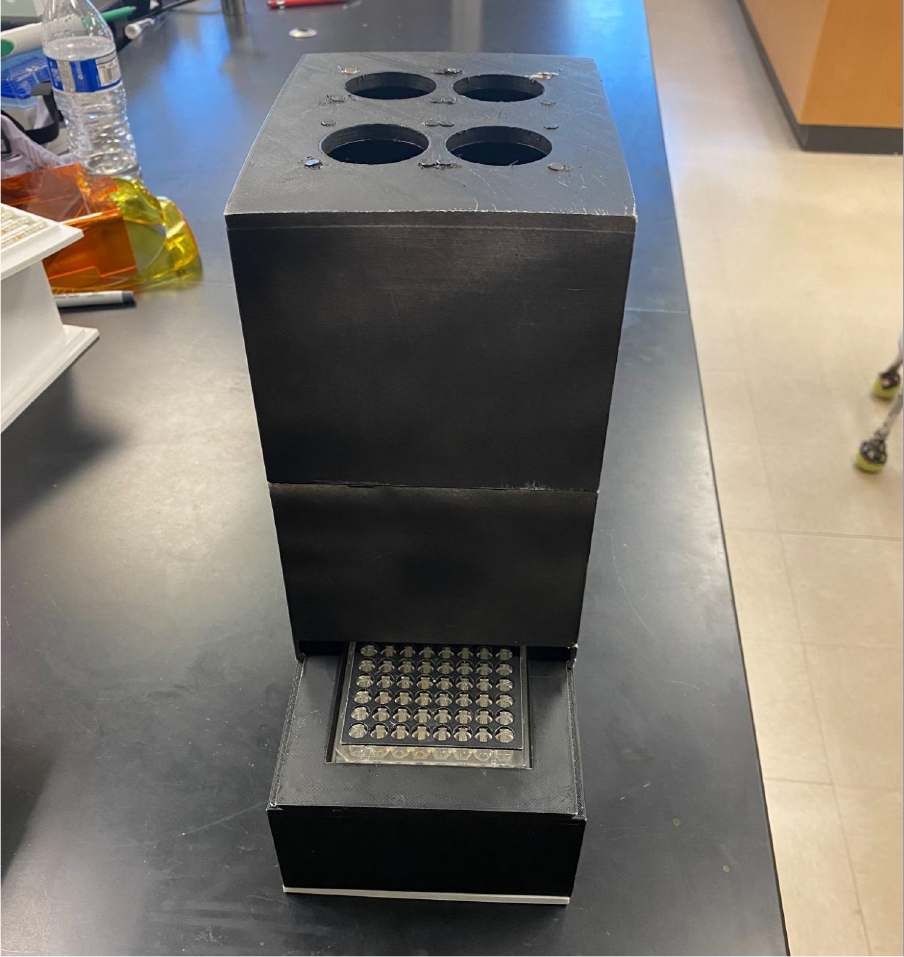
Image of 3D printed plate reader with well plate inserted.

**Figure 6.**
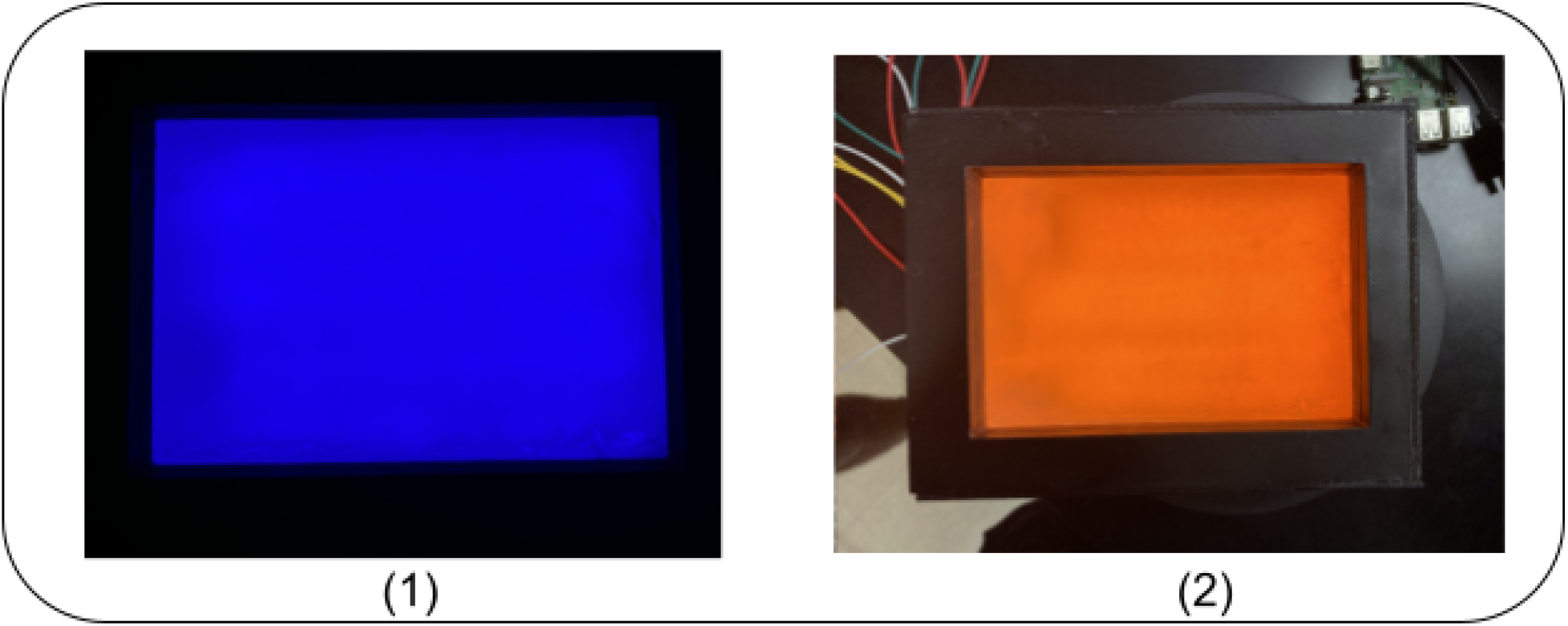
6-1. Even light distribution of the 440 nm excitation light. 6-2. Even light distribution of the 600nm optical density light.

**Figure 7.**
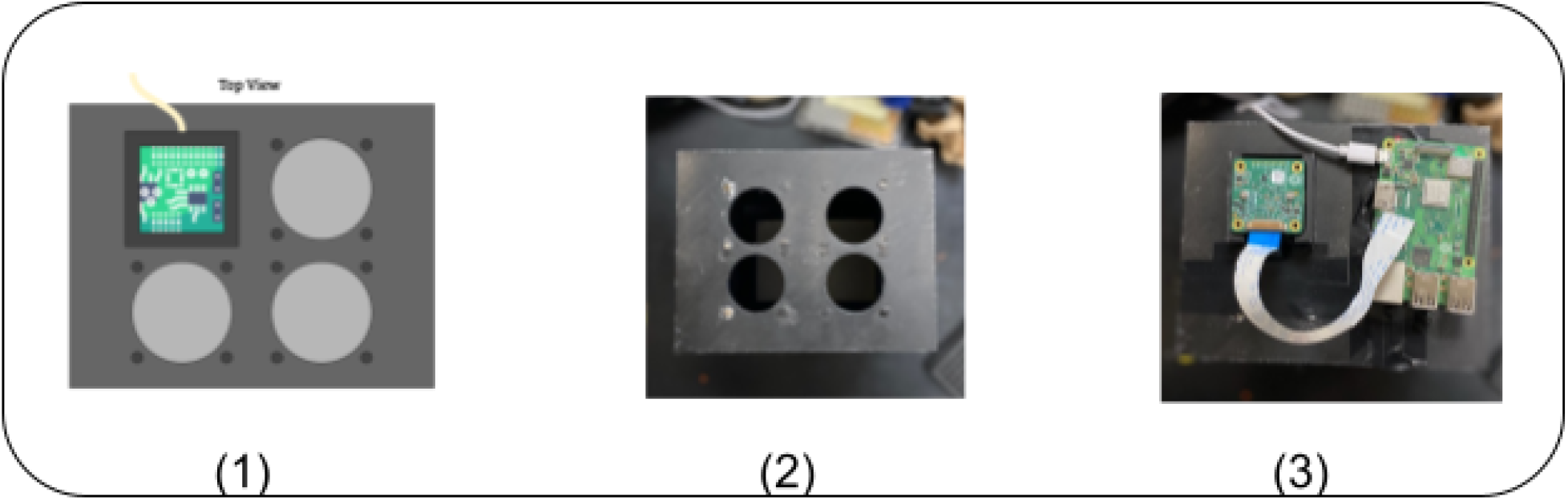
7-1. Schematic of camera movement over microplate. 7-2. 3D printed top cover with holes for camera movement. 7-3. Top cover with camera and raspberry pi attached

## Problems and Solutions

### Camera Movement and Perspective Adjustion

A common problem with camera-based devices is lens distortion and camera perspective [1]. In order to counteract lens distortion, we utilized a 16mm focal length lens which minimizes the fisheye effect compared to other lenses. On the other hand, we minimized the camera perspective problem by moving the camera over multiple positions of the well plate (Figures. 6) and utilizing a perspective transformation algorithm.

### Well Plate Cover

Another common problem with extracting luminescence from a clear well plate image is signal distortion [1]. To solve this problem we developed a well plate cover that fits into most 96 holed well plates (Fig. 8). With this, the signal from one well won’t affect the readings of the surrounding wells.

**Figure 8.**
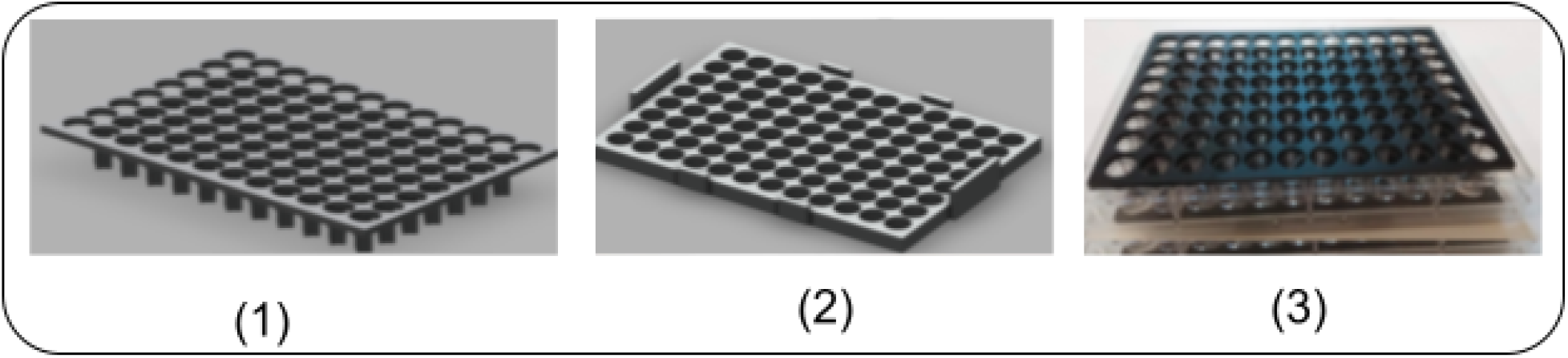
8-1. CAD of top cover. 8-2. CAD of bottom cover. 8-3. 3D printed well plate cover attached to a well plate

### Camera Settings

In order to get the most precision from the Raspi camera in a low-light setting, we adjusted various camera settings such as ISO, exposure, shutter speed, etc., to manipulate the output image.

Below are the camera settings used:

- Shutter speed: 1000 ms
- ISO: 800
- Exposure: Night mode (this is built into the raspi still library on the Raspberry Pi) These camera settings are ideal for photography of low-light scenes and worked well for capturing well plates for Plate-Q [7].

## Software Processing

### Overview

Plate-Q relies on a standardized Raspberry Pi HQ camera for the quantification of fluorescence and optical density, which is done through a software pipeline (Figure 9) that extracts the image features and maps those values to laboratory-grade plate reader values. The pipeline starts by taking pictures in triplicates at 4 different positions across the well plate. Each image is then converted into grayscale and applied a perspective transformation to analyze the perceived brightness. Then to isolate the wells in the image, Otsu’s Thresholding is used. Then a gamma correction is applied to the image to encode it into a linear luminescence scale. Finally, the brightness values of the triplicates are extracted and averaged to eliminate any outliers (Figure 10). Then the final value is stored in a data frame (table) that is passed through a mathematical regression model to output laboratory-grade plate reader values.

**Figure 9.**
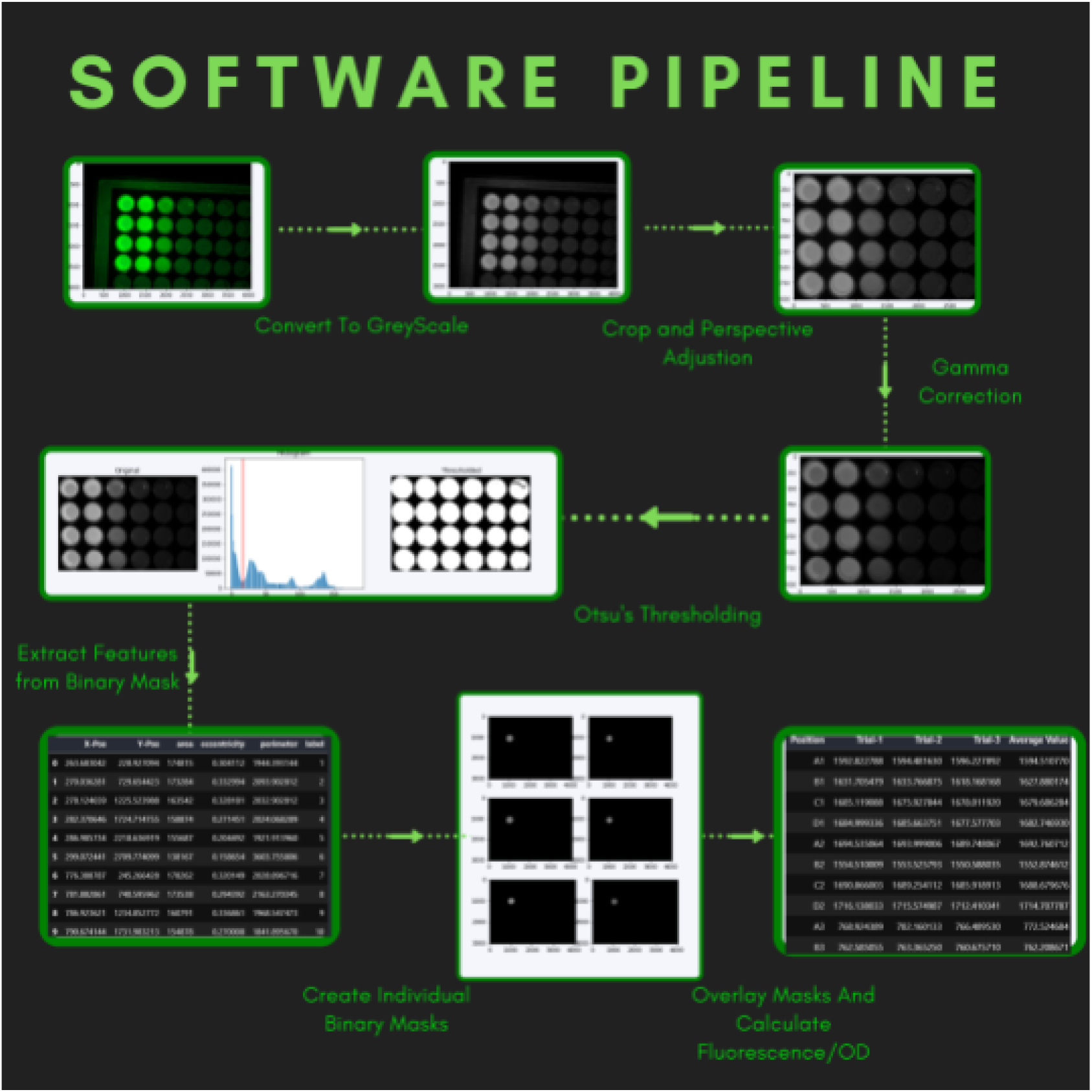
Visual representation of software pipeline.

**Figure 10.**
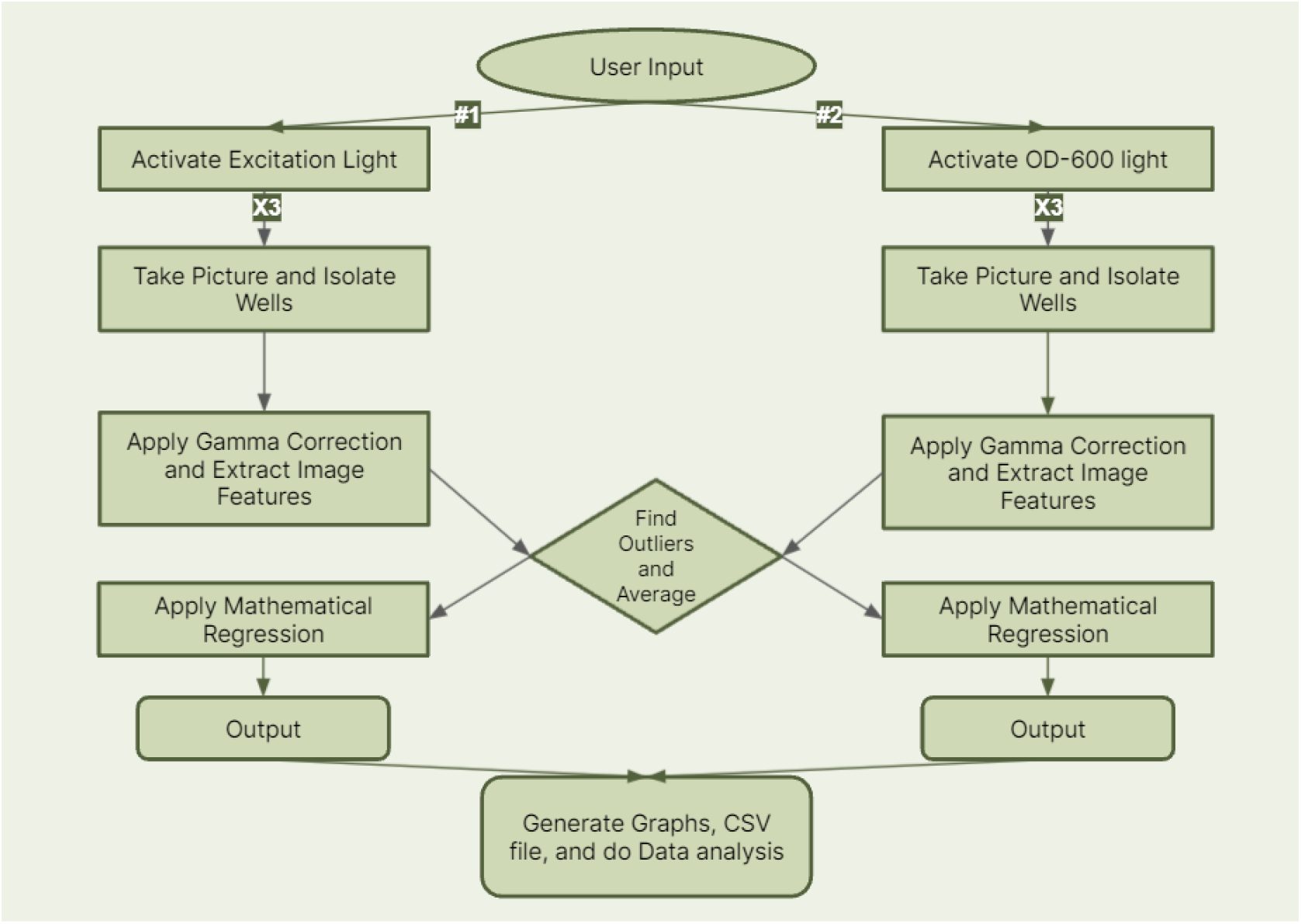
Diagram of Plate-Q software pipeline.

### Perspective Transformation Algorithm

To solve the problem of camera perspective, we utilized the perspective transformation algorithm. This type of transformation does not preserve parallelism, length, and angle, but instead preserves collinearity and incidence. This means that the straight lines will remain straight even after the transformation. The perspective transformation can be represented by the formula (1) shown below, where,(x, y) are the input points,(x’, y’) are the transformed points, and M is the 3*3 transformed Matrix.

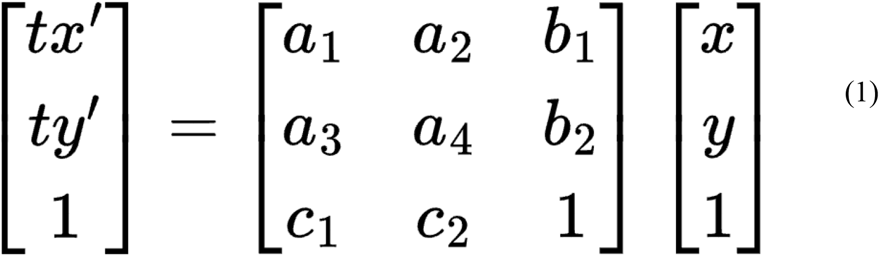

Matrix M is a combination of an image transformation matrix(a1-a4), translation vector(b1-b2), and projection vector(c1-c2). With Matrix M having 8 degrees of freedom we are able to select 4 points in the input image and map these 4 points to the desired locations in the output image. In our case, we are able to identify the corners of interest in the image and calculate the perspective transformed image. With this, we are able to minimize the camera perspective problem.

### Otsu’s Method and Well Isolation

Otsu’s Method, a computer vision technique, is used to isolate wells in an image (Figure 11). Otsu’s method[2] is a variance-based technique to find the threshold value where the variance between the foreground and background pixels is the least. With this, the algorithm iteratively applies different values of thresholds to find the value that minimizes the within-class variance between the two classes. The formula to find the within-class variance at any threshold can be represented by the equation (2) where t represents the threshold value, ωbg and ωfg represent the probability of the number of pixels for each class at a threshold t, and σ2 represents the color values [3].

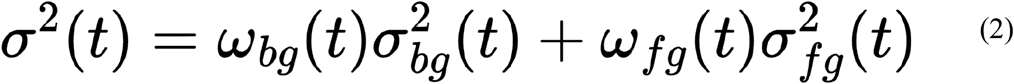

**Figure 11.**
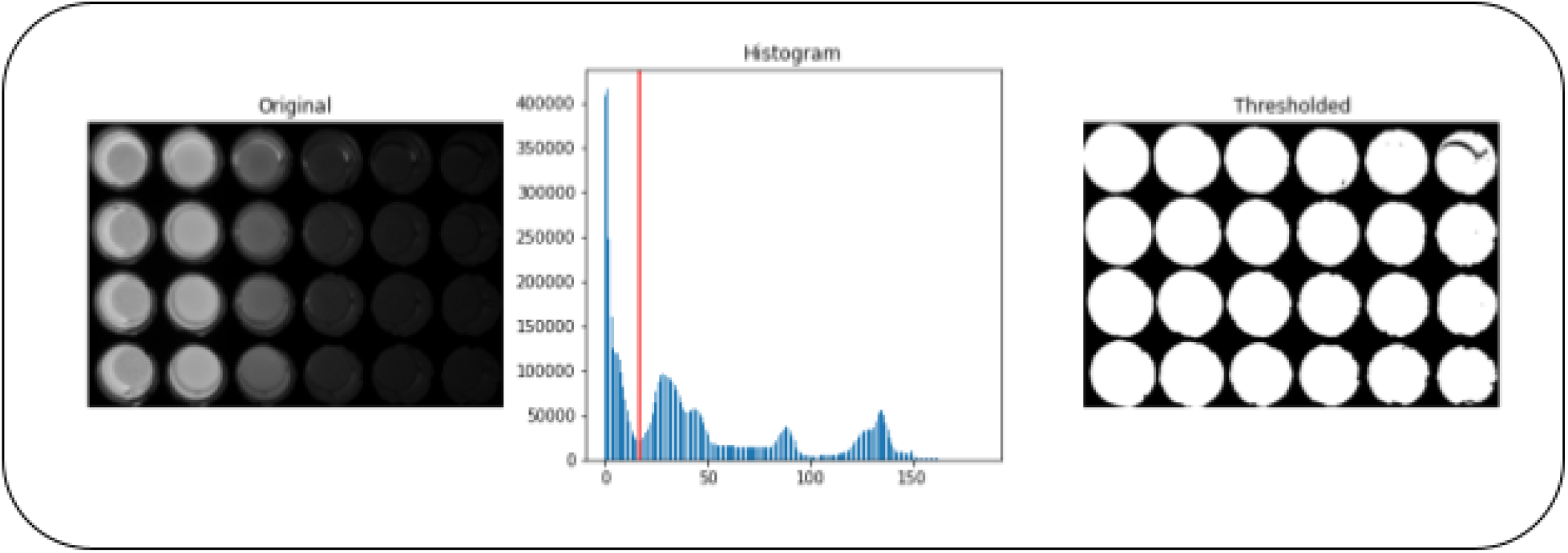
Example of Otsu’s Method applied to fluorescence data.

**Figure 12.**
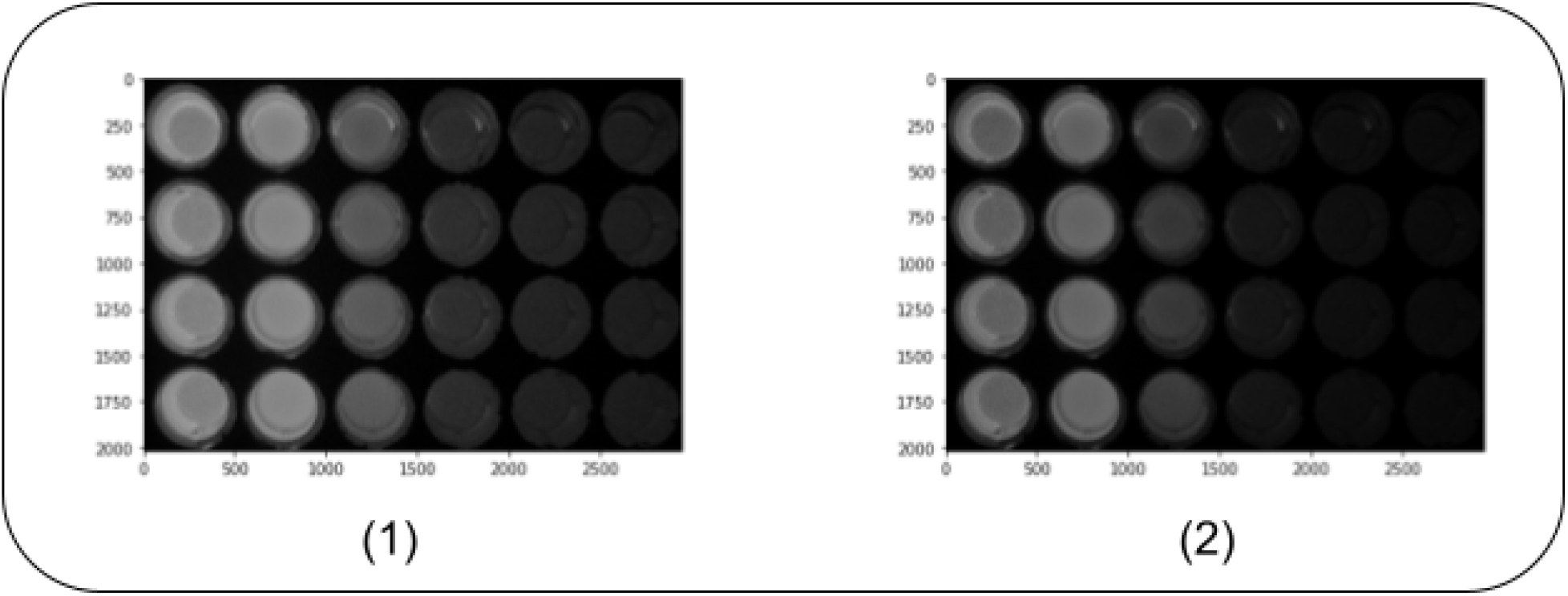
12-1. Image before gamma correction. 12-2. Image after gamma correction.

For our situation, our image is in grayscale so pixel values are between 0-255. After iteration and finding that the threshold that minimizes the within-class variance is *x*, all pixel values greater than or equal to *x* become the foreground of the image while all other pixels become the background. With this, we are able to isolate the wells from the background as the well’s pixels are brighter than the background of the image.

### Gamma Correction

Gamma correction is a process of translating the brightness measured by camera sensors to the perceived brightness of human eyes. It is used to compensate for the non-linear relationship between luminescence and brightness as measured by the camera (Figure 11). Pictures taken by a camera are gamma encoded in an sRGB format, which records red, green, and blue light ranges in a non-linear method. Gamma correction applies the inverse of this encoding function to transform the non-linear sRGB [4].

To apply a gamma correction to an input image, each pixel value from a scale of 0 to 255 is converted to a scale of 0 to 1. The following function (3) is applied to the pixel values of the input image: where I is the input image, G is the gamma value, and O is the output image.

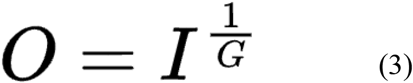

### Calculating Fluorescence and Optical Density

In order to calculate fluorescence using brightness values extracted from an image, the following equation is used (4) where I_0_ is the amount of light that initially hits the sample, T is the amount of light that transmits through the sample, and OD is the calculated optical density. I_0_ is measured by scanning a clear plate or a blank well through Plate-Q and extracting the 0 to 255 pixel brightness value, and T is measured by scanning a filled well through Plate-Q and extracting the brightness value.

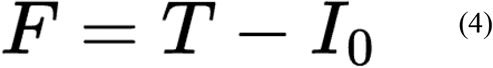

In order to calculate the optical density using brightness values extracted from an image, the following equation is used (5) where I_0_ is the amount of light that initially hits the sample, T is the amount of light that transmits through the sample, and OD is the calculated optical density. I_0_ is measured by scanning a clear plate or a blank well through Plate-Q and extracting the 0 to 255 pixel brightness value, and T is measured by scanning a filled well through Plate-Q and extracting the brightness value [5].

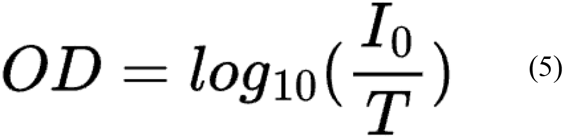

### Regression

To find the optimal gamma correction value and to map values to real plate reader values, we applied regression comparing calculated values of Plate-Q with a wide range of gamma correction applied to real plate reader values. To find the best gamma correction value, we tried to linearize the data by applying a wide range of gamma values. When iterating through a wide range of gamma correction values and applying linear regression on the data, we looked for the gamma value that would result in the r^2^ (correlation coefficient) value closest to 1. In addition, the resulting linear regression equation was used to map the values to a laboratory-grade plate reader. Without analysis we were able to determine the optimal gamma correction value of Fluorescence to be .7 with a regression equation of *y* = 31.1 + 9. 8 *(raw PlateQ Value)* (Figures. 13 & 14). On the other hand, optical density’s optical density value was .23 with a regression equation of *y* = . 0635 + 1. 5(*raw PlateQ Value*).

**Figure 13.**
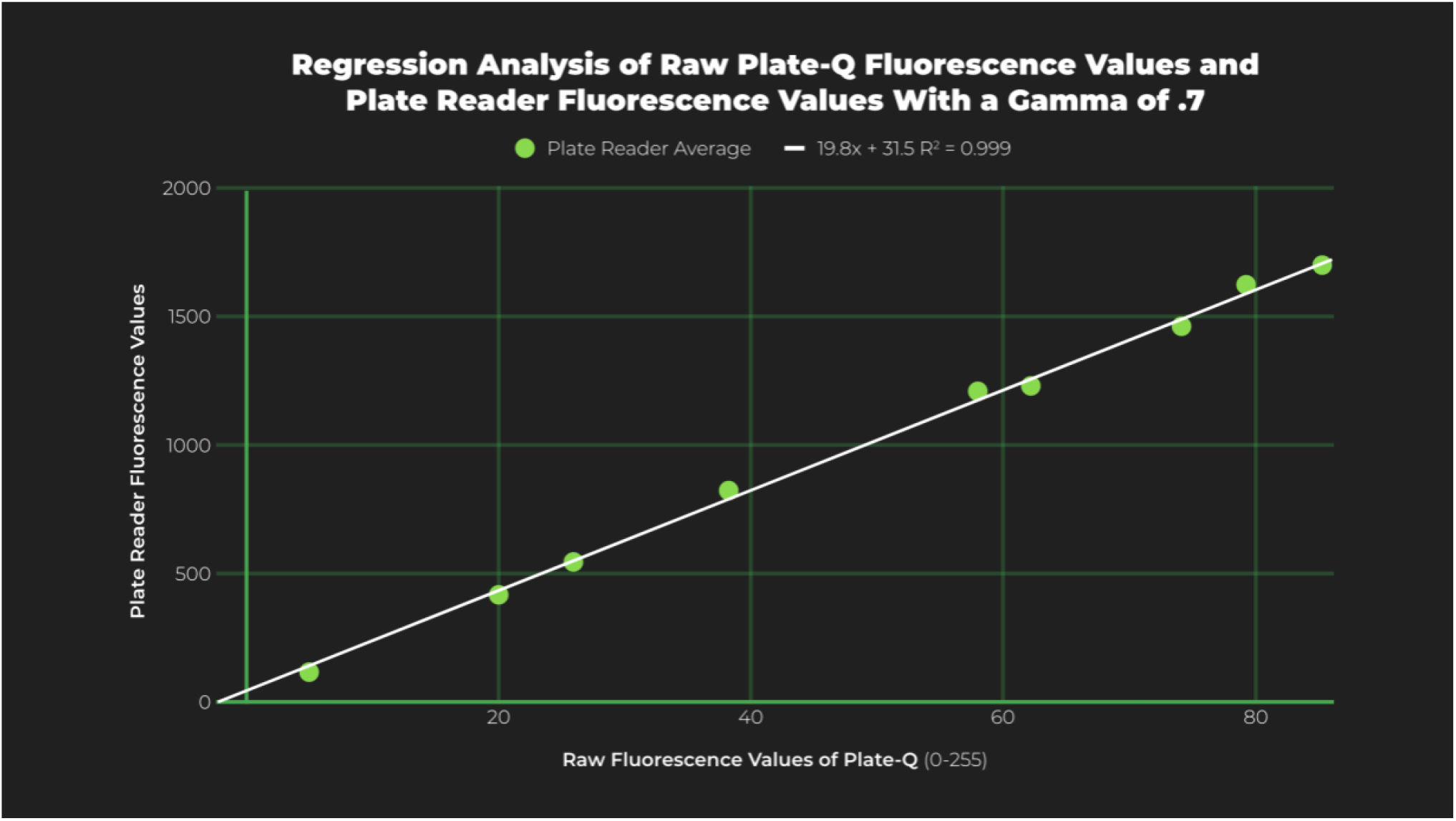
Image of linear regression on raw OD values from Plate-Q.

**Figure 14.**
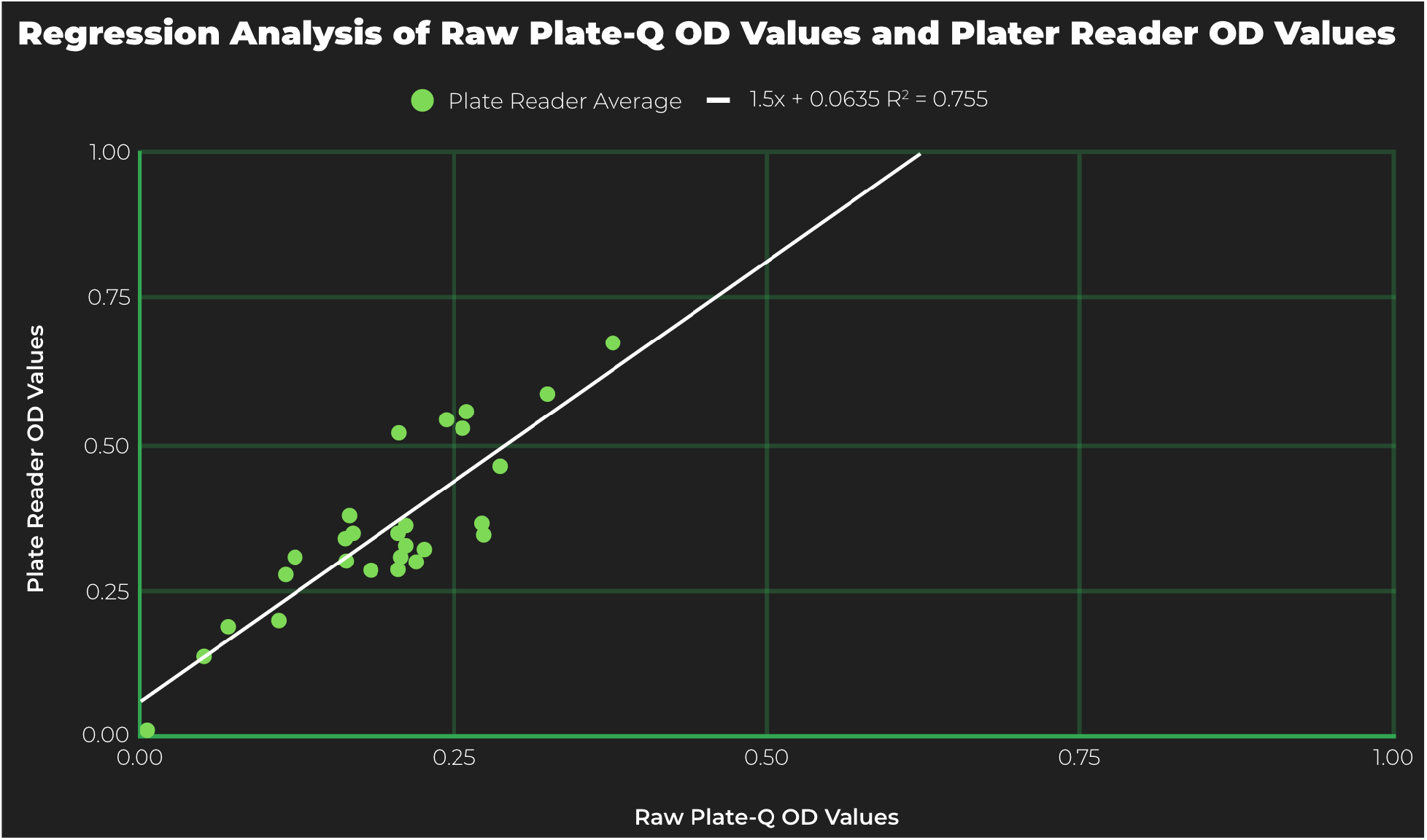
Image of linear regression on raw OD values from Plate-Q.

## Results

After obtaining results, we analyzed fluorescence values and determined that Plate-Q works best at greater brightness values between 400 to 2000 with a percent error of 4.7% (Figure. 15). Because Plate-Q overestimates for lower fluorescence values, this shows that the camera sensor has trouble quantifying lower brightness values.

**Figure 15.**
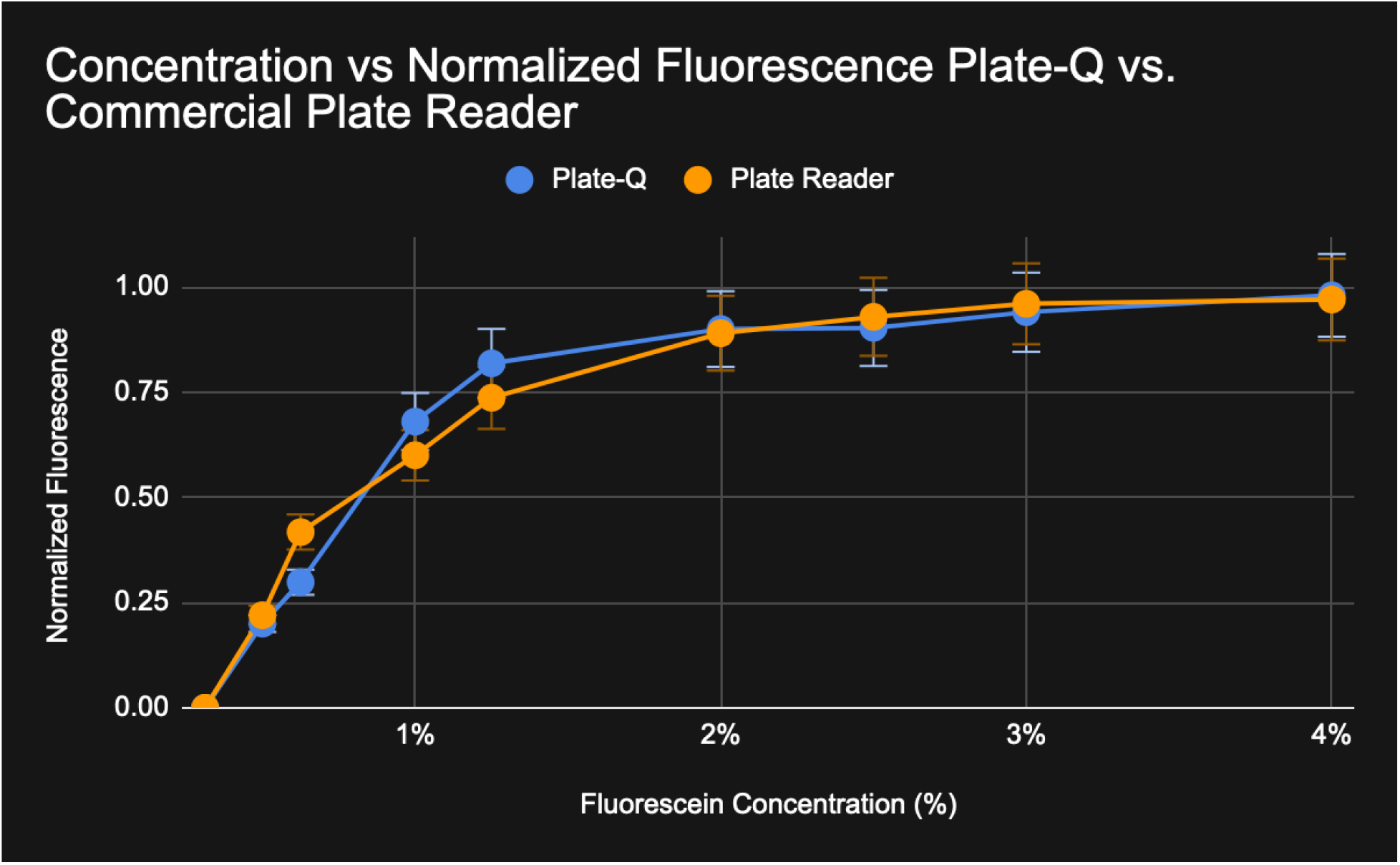
Comparison of fluorescence in commercial microplate reader and Plate-Q at different concentrations and ranges.

Similar to fluorescence measurement, optical density (OD) in Plate-Q performs more accurately at higher brightness values than lower brightness values. In comparison with a laboratory microplate reader, Plate-Q tends to overestimate optical density (OD) at values approximately greater than 0.3. This is likely due to the low sensitivity of the Raspberry Pi camera sensor at low light settings, because higher OD values are associated with less light transmission through a sample, or lower brightness values. Due to this, the returned output of Plate-Q will be less reliable at a higher OD. Plate-Q was able to output an average percent error of 18.74% (Figure 16.)

**Figure 16.**
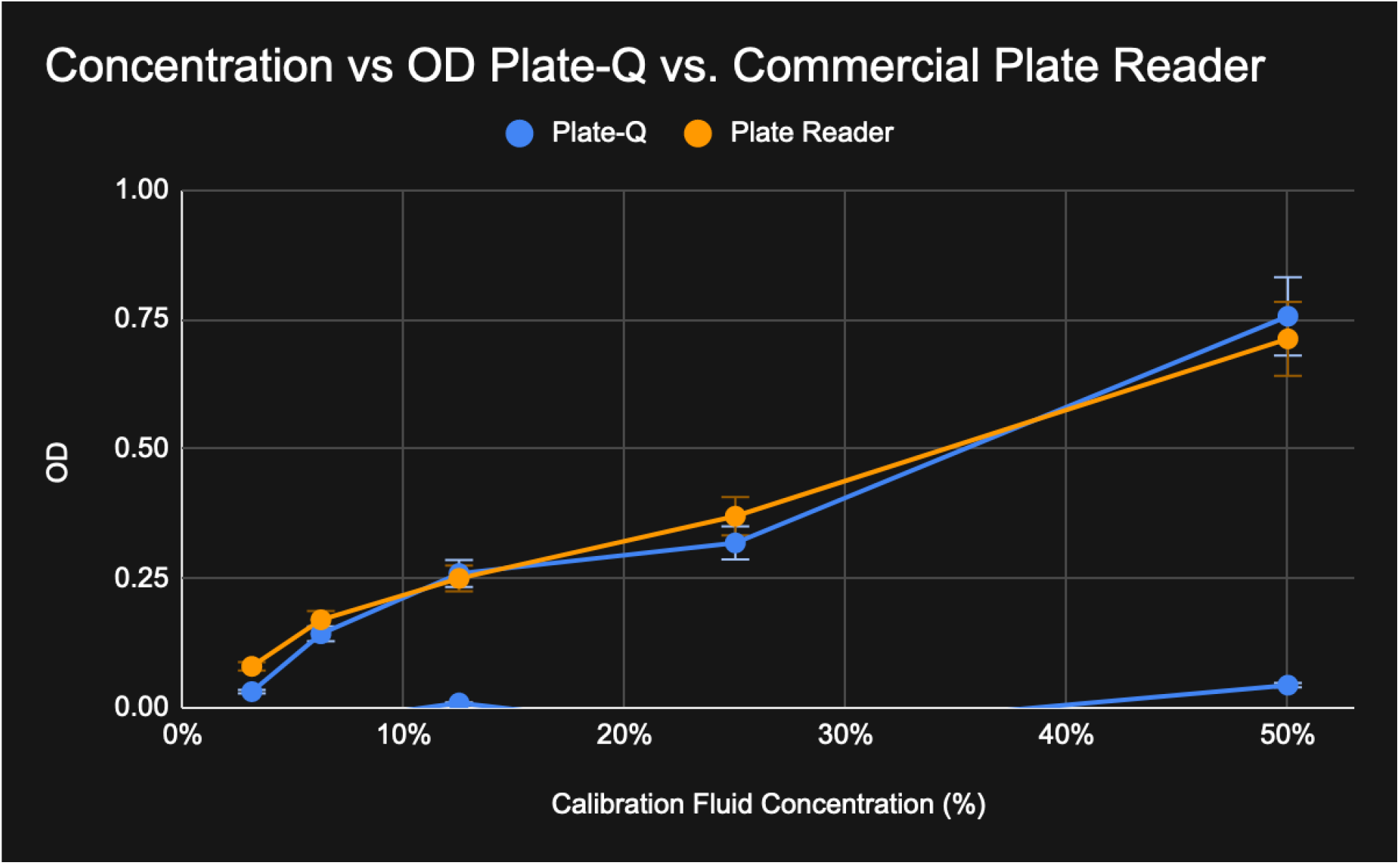
Comparison of optical density output between Plate-Q and a laboratory microplate reader.

Overall, Plate-Q was able to have less variability in data, but had trouble with lower brightness values. Moving forward, we plan on testing different shutter speed settings to manipulate the camera sensitivity to brightness in addition to using more sensitive camera sensors.

## Discussion

### Summary

In typical labs, microplate readers are very helpful in quantifying green fluorescence protein (GFP) and Optical Density (OD) from a 96-well standard plate; however, laboratory-grade plate readers can cost up to $20,000 which is cost-prohibitive for underfunded labs. To address this issue, we created Plate-Q: a frugal and open-source microplate reader. At just under $150, Plate-Q is an inexpensive way to quantify both fluorescence and optical density (OD) of samples. Rather than using optical sensors found in laboratory-grade plate readers[6], Plate-Q takes advantage of a Raspberry Pi camera to capture images of a well plate and extracts image features using computer vision and machine learning algorithms. A 440 nm wavelength excitation light is used to measure GFP fluorescence using a 510 nm filter for emission, and a 600 nm light source is used for OD without any emission filter. Plate-Q is completely open-source and users can customize the design to scan for other fluorescent proteins by replacing the light source and filter for different wavelengths. Users can retrain the Plate-Q algorithm to take measurements affordably and sustainably.

### Future Development

With the proven success of Plate-Q for fluorescence and optical density, we plan to develop two future versions to target different demographics. The first of which will be a more frugal version that utilizes the mobile phone camera as the quantification sensor. But with the use of a mobile phone camera, there will be variability between different devices due to different hardware and software packages. To counter this we plan on developing a mobile app for quantification that will allow users to calibrate their camera to get the optimal settings for use. The other version will build off of Plate-Q and address the problem of lower brightness quantification. We plan on utilizing the camera sensors from popular astrophotography cameras to develop a custom camera that will have higher sensitivity at low light settings. In addition, we plan on adding functionalities for luminescence, time-resolved fluorescence, and colorimetry to provide functionalities of laboratory-grade microplate readers. With this, we hope to provide a low-cost alternative that will allow underfunded labs and developing countries to perform biotechnology research that can similarly be achieved by its laboratory-grade counterparts.

